# Lower Aperiodic EEG Activity is Associated with Reduced Verbal Fluency Performance Across Adulthood

**DOI:** 10.1101/2024.08.13.607827

**Authors:** Daniel J McKeown, Emily Roberts, Anna J. Finley, Nicholas J. Kelley, Hannah A.D. Keage, Victor R Schinazi, Oliver Baumann, Ahmed A Moustafa, Douglas J Angus

## Abstract

Age-related cognitive decline associations with human electroencephalography (EEG) have previously focused on periodic activity. However, EEG primarily consists of non-oscillatory aperiodic activity, characterised with an exponent and offset value. In a secondary analysis of a cohort of 111 healthy participants aged 17 – 71 years, we examined the associations of the aperiodic exponent and offset in resting EEG with a battery of cognitive tests consisting of the Colour-Word Interference Test, Wechsler Adult Intelligence Scale IV Digit Span Test, Rey Auditory Learning Test, Delis-Kaplan Executive Function System Trail Making Test, and the Verbal Fluency Test. Using Principal Component Analysis and K-Means Clustering, we identified clusters of electrodes that exhibited similar aperiodic exponent and offset activity during resting-state eyes-closed EEG. Robust linear models were then used to model how aperiodic activity interacted with age and their associations with performance during each cognitive test. Offset by age interactions were identified for the Verbal Fluency Test, where smaller offsets were associated with poorer performance in adults as early as 33 years of age. Greater aperiodic activity is increasingly related to better verbal fluency performance with age in adulthood.

## INTRODUCTION

Investigations into the electroencephalogram (EEG) have historically focused on oscillatory activity across the power spectrum. However, EEG predominantly consists of non-oscillatory (i.e., aperiodic) 1/*f*-like patterns of activity (He, 2014). Once considered artifactual noise, aperiodic activity is of neurophysiological significance and functionally important (Brake et al., 2024; Donoghue, Haller, et al., 2020). The exponent (the slope of the EEG broadband) and offset (*y*-intercept of the EEG broadband) provide an indication of excitation: inhibition (E:I) balance and neural spiking rate, respectively (Donoghue, Haller, et al., 2020; Gao et al., 2017). Dynamic changes in the aperiodic activity of EEG can be due variation in the input of various neural generators, integrity of the signal-to-noise ratio, and task demand (Donoghue et al., 2022; Gao et al., 2017; Gao et al., 2020; Miller et al., 2014; Miller et al., 2009). Variation in the exponent and offset are predictors of task performance (Hohn et al., 2024; Immink et al., 2021; Pathania et al., 2021) and age-related cognitive decline (Finley et al., 2024; Tran et al., 2020; Voytek et al., 2015), and can be considered biomarkers of brain disorders (Arnett et al., 2022; McKeown et al., 2023).

Aging is associated with wide-scale changes in aperiodic activity reflecting non-pathological changes in neuronal networks (Finley et al., 2024; Voytek et al., 2015). The exponent reduces with age, suggesting increases in E:I balance, while the offset declines, likely reflecting reduced neural firing (Clark et al., 2024; Finley et al., 2024; Merkin et al., 2023; Voytek et al., 2015). Recently, this has been documented to occur as early as 4 – 12 years of age (Hill et al., 2022). Aperiodic activity also plays a role in maintaining cognitive function throughout life (Pathania et al., 2022). Previous research shows that the exponent and offset are negatively correlated with reaction time (Euler et al., 2024), perceptual sensitivity (Immink et al., 2021), processing speed (Ouyang et al., 2020), and selective attention performance (Waschke et al., 2021). However, these associations do not appear across all ages or tasks, suggesting that the relationship between aperiodic activity and cognition varies with age and task (Cesnaite et al., 2023; Pei et al., 2023; Waschke et al., 2021).

Recently, Euler et al. (2024) found no association between the resting EEG aperiodic exponent and individual constructs of working memory, perceptual reasoning, processing speed, and verbal comprehension from the Wechsler Adult Intelligence Scale (WAIS) subtests in 166 participants (18 – 52 years). However, combining the eight WAIS subtests revealed that higher exponents were associated with greater general ability. While Cesnaite et al. (2023) found no associations between aperiodic activity and cognitive performance in an older cohort (n = 1703, 60 – 80 years), Smith et al. (2023) linked higher exponents with better general cognitive function but not age (n = 86, 50 – 80 years). While these effects may vary by age and task, most studies focus on limited age ranges, excluding early-life (Mage = 54.86, 36 - 83 years; Finley et al., 2024), and including exclusively older cohorts (50 - 80 years; Cesnaite et al., 2023; Smith et al., 2023). To date, no studies have examined associations between aperiodic activity, and cognitive ability inclusively across early, mid, and late life.

We present a secondary exploratory analysis of resting-state EEG from the Stimulus-Selective Response Modulation (SRM) project (Hatlestad-Hall et al., 2022; Rygvold et al., 2022). This dataset consists of resting-state EEG and performance measures during a battery of neuropsychological tests from 111 healthy participants aged 17 – 71 years. Aperiodic activity was derived from resting EEG and cognitive performance was derived from performance measures of a cognitive test battery consisting of a Colour-Word Interference Test, Wechsler Adult Intelligence Scale IV Digit Span Test, Rey Auditory Learning Test, Delis-Kaplan Executive Function System Trail Making Test, and a Verbal Fluency Test. This dataset was chosen as the cognitive test battery covers a broad range of cognitive processes. We hypothesise that lower aperiodic activity will be associated with lower scores in each cognitive test and that these associations will strengthen as age increases.

## MATERIALS AND METHODS

### Code Availability

The processes and properties of the original data are described in full by Rygvold et al. (2021) and Hatlestad-Hall et al. (2022), with relevant details for the current study described below. Processed data and scripts for the current study are available at https://github.com/MindSpaceLab/Aperiodic_SRM_Cognition.

### Participants

The original sample of 111 participants is available at OpenNeuro (“OpenNeuro Dataset ds003775 (SRM Resting-state EEG)”; https://openneuro.org/datasets/ds003775/versions/1.2.1; Hatlestad-Hall et al., 2022). Participants were aged between 17 and 71 (mean age = 37.5, SD = 13.99; Figure 1) and consisted of 68 females. All participants were neurologically and psychiatrically healthy individuals (self-reported) and were unmedicated. The Bond University Human Research Ethics Committee approved our secondary analysis of this data (approval number: DA202221102).

**Figure 1.**
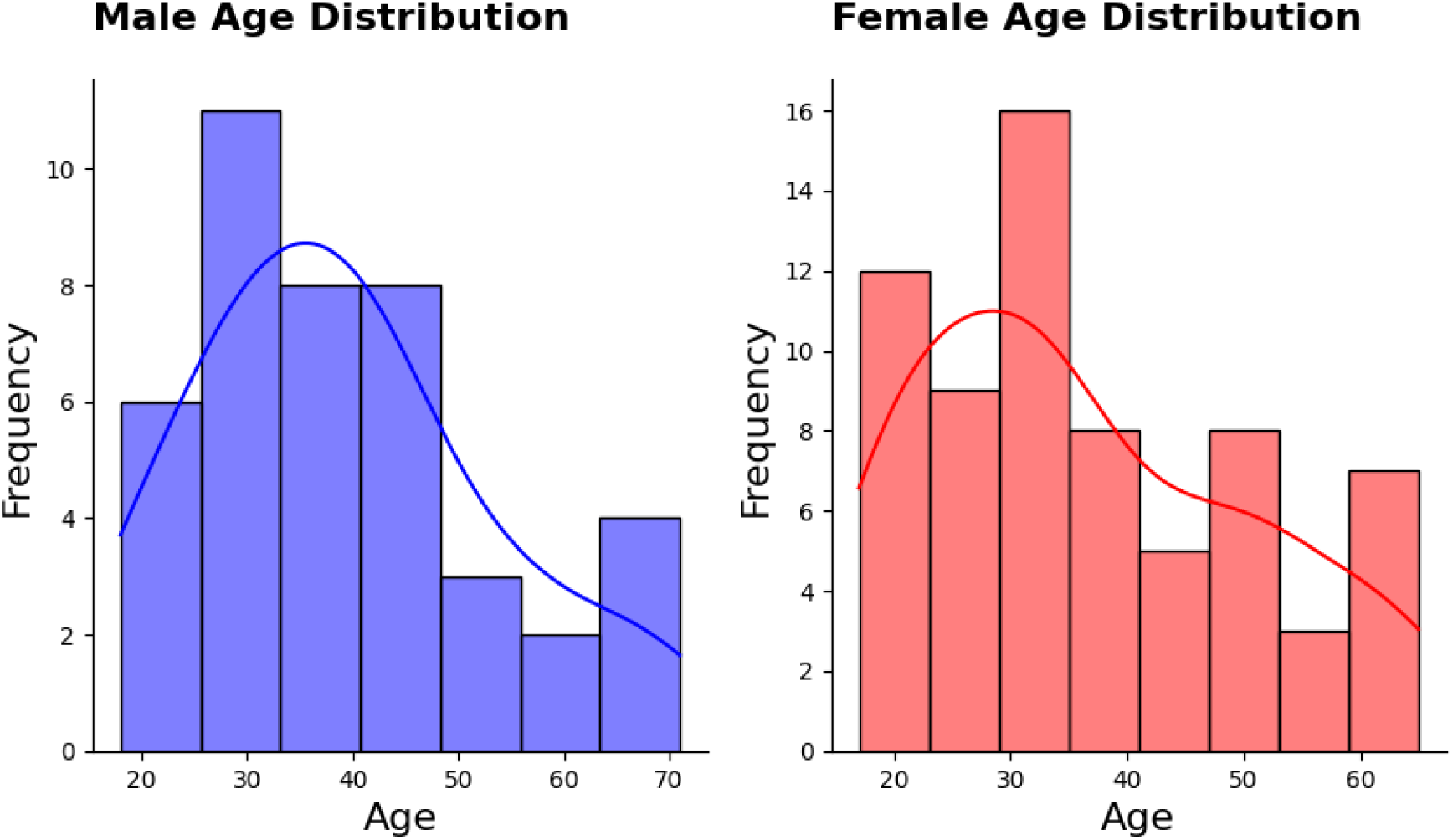
Age distribution for male and female participants.

### EEG Acquisition

Full details on the experimental procedure can be found at Hatlestad-Hall et al. (2022) but are summarised here. Four minutes of resting-state eyes-closed EEG data was collected towards the end of the experimental protocol, approximately 45 minutes into the session after the battery of neuropsychological tests using a 64-channel (Ag-AgCl electrodes) BioSemi ActiveTwo system (BioSemi B.V., Amsterdam) arranged according to the extended 10-20 system (10-10). EEG data was initially acquired with a sampling rate of 1024 Hz without online filtering. Four external electrodes were used to detect eye movement. These consisted of lateral, and inferior/superior to the right eye (LO1, LO2, IO1, and SO2). Two electrodes were positioned on each earlobe for grounding and referencing of the signal (locations A1 and A2).

### EEG Preprocessing

The current study utilised the cleaned raw EEG data files provided by Hatlestad-Hall et al. (2022). The initial preprocessing procedure consisted of re-referencing to the average reference and an off-line anti-aliasing filter applied, algorithmically detecting bad segments and channels then removing them from the data, applying a high-pass filter of 1 Hz, using ZapLine to remove power line noise, calculating independent components with the second order blind identification (SOBI) algorithm, implementing ICLabel (Delorme et al., 2024; Pion-Tonachini et al., 2019) to subtract from the data the components that were considered eye or muscle artifact with 85% certainty, and applying a low-pass filter of 45 Hz. A ‘strict’ data rejection procedure was implemented and as a result, 64.1% of the EEG data files retained > 90% of their channels, while 23.5% retained 75-90% of their channels. Channels considered to be of poor quality were interpolated. After preprocessing, continuous data was segmented into 4000 ms epochs with no overlap.

Our secondary analysis was conducted in Python (v3.11.5). Our secondary analysis begun by averaging the epochs to generate a single 4000 ms segment for calculation of power spectral densities (PSD). An average of 54 ± 3.37 epochs (range = 22 – 55) were included in PSD calculations. We used a Fast Fourier Transformation (FFT; 2000 ms Hamming window, 50% overlap) to transform the data into the PSD from 1 Hz to the Nyquist max. The aperiodic exponent and offset were quantified from the PSD using the *Specparam* package (https://github.com/fooof-tools/fooof; Donoghue, Haller, et al., 2020). For each PSD we fit the *Specparam* model between 1 and 40 Hz (peak width limits: 1.6 – 6; max number of peaks: 8; minimum peak height: .05; peak threshold: 1.5 SD, aperiodic mode: fixed) with a .25 Hz frequency resolution. Aperiodic exponent and offset, as well as the parameterised periodic activity in the alpha band (8 – 13 Hz) were extracted. Fitting of the *Specparam* model performed excellently (mean *R*^2^ across the scalp and across all participants = .98 ± .005). No participants had fits < .9 *R*^2^ in more than 50% of their channels nor extreme exponent or offset values (> 3 SD). Retainment of parameterised alpha power was also excellent with 94.95% of channels having retained a parameterised peak in the alpha band.

### Cognitive Test Battery

Participants in the study completed a battery of neuropsychological tests consisting of: the Colour-Word Interference Test from the Delis-Kaplan Executive Function System (D-KEFS), the Digit Span Test from the Wechsler Adult Intelligence Scale-IV, the Rey Auditory Verbal Learning Test, the Trail Making Test from the D-KEFS, and the Verbal Fluency Test from the D-KEFS. Details of the variables and the participant performance of each test can be found in Hatlestad-Hall et al. (2022). Outcomes measures for the tests consisted of time in seconds to complete the tasks for the Colour-Word Interference Test and the Trail Making Test, and total score for the Digit Span Test, Rey Auditory Learning Test and Verbal Learning Test. A description of each neuropsychological test used in the original study by Hatlestad-Hall et al. (2022) is detailed below, and the performance of the participants for each cognitive test is presented in Table 1.

**Table 1:** Performance scores from each neuropsychological test of the cognitive test battery.

| Test | Variable | Mean ± SD | Range |
| --- | --- | --- | --- |
| RAVLT | ravlt_1 | 8.4 ± 2.2 | 4 – 14 |
| | ravlt_5 | 13.9 ± 1.3 | 10 – 15 |
| | ravlt_tot | 60.5 ± 7.6 | 43 – 74 |
| | ravlt_imm | 12.9 ± 2.3 | 5 – 15 |
| | ravlt_del | 13.0 ± 2.3 | 5 – 17 |
| | ravlt_rec | 14.4 ± 1.6 | 12 – 15 |
| | ravlt_fp | 0.3 ± 0.8 | 0 – 7 |
| WAIS-IV DS | ds_forw | 9.6 ± 2.2 | 5 – 15 |
| | ds_back | 9.1 ± 2.0 | 4 – 16 |
| | ds_seq | 9.5 ± 2.2 | 5 – 15 |
| | ds_tot | 28.2 ± 5.0 | 17 – 44 |
| TMT | tmt_2 | 28.3 ± 11.4 | 13 – 75 |
| | tmt_3 | 27.4 ± 12.9 | 11 – 77 |
| | tmt_4 | 70.3 ± 30.4 | 29 – 202 |
| CWIT | cw_1 | 29.1 ± 5.7 | 20 – 53 |
| | cw_2 | 21.0 ± 3.4 | 14 – 30 |
| | cw_3 | 49.4 ± 12.5 | 31 – 100 |
| | cw_4 | 56.7 ± 14.9 | 37 – 130 |
| VF | vf_1 | 47.8 ± 12.5 | 19 – 84 |
| | vf_2 | 51.2 ± 11.6 | 14 – 88 |
| | vf_3 | 16.1 ± 3.3 | 7 – 24 |

*CWIT, Colour-Word Interference Test; WAIS-IV DST, Wechsler Adult Intelligence Scale IV Digit Span Test; RAVLT, Rey's Adult Verbal Learning Test; DKEFS TMT, Delis-Kaplan Executive Function System Trail Making Test; VFT, Verbal Fluency Test. Description of each variable can be found in the Cognitive Test Battery section of the Materials and Methods. Adapted from Hatlestad-Hall et al. (2022).*

#### Delis-Kaplan Executive Function System Colour-Word Interference Test

The D-KEFS Colour-Word Interference Test was used to assess reading speed and executive functioning capabilities. Scores were quantified as follows: 1) number of seconds elapsed before completion of the test’s colour only condition (cw_1), 2) number of seconds elapsed before completion of the test’s reading only condition (cw_2), 3) number of seconds elapsed before completion of the test’s interference condition (cw_3), 4) number of seconds elapsed before completion of the test’s interference and switching condition (cw_4; Hatlestad-Hall et al., 2022). In our secondary analysis, the average score for cw_1, cw_2, cw_3, and cw_4 was quantified to represent Colour-Word Interference Test performance.

#### Wechsler Adult Intelligence Scale-IV Digit Span Test

The Digit Span Test was used to assess attentional performance. Scores were quantified as: 1) achieved score in the test’s forward condition (ds_forw), 2) achieved score in the test’s backward condition (ds_back), achieved score in the test’s sequencing condition (ds_seq), and total score across all conditions (ds_tot; Hatlestad-Hall et al., 2022). In our secondary analysis we used ds_tot to quantify Digit Span Test performance.

#### Rey Auditory Verbal Learning Test

The Rey Auditory Verbal Learning Test was used to assess verbal memory performance. Scores were quantified as: 1) number of items correctly recalled after first learning trial (ravlt_1), 2) number of items correctly recalled after fifth learning trial (ravlt_5), 3) number of items correctly recalled across all five learning trials (ravlt_tot), 4) number of items correctly recalled immediately after the final learning trial (ravlt_imm), 5) number of items correctly recalled 30 minutes after the final learning trial (ravlt_del), 6) number of items correctly recognised from a list (ravlt_rec), 7) number of false positive responses during the recognition task (ravlt_fp; Hatlestad-Hall et al., 2022). In our secondary analysis we calculated the average of ravlt_tot, ravlt_imm, and ravlt_del to quantify Verbal Learning Test performance.

#### Delis-Kaplan Executive Function System Trail Making Test

The D-KEFS Trail Making Test was used to quantify processing speed and executive functioning capabilities. Performance was quantified by: 1) number of seconds elapsed before completion of the test’s number condition (tmt_2), 2) number of seconds elapsed before completion of the test’s letter condition (tmt_3), and 3) number of seconds elapsed before completion of the test’s switching condition (tmt_4; Hatlestad-Hall et al., 2022). In our secondary analysis we quantified Trail Making Test performance by calculating the average of tmt_2, tmt_3, and tmt_4.

#### Delis-Kaplan Executive Function System Verbal Fluency Test

The D-KEFS Verbal Fluency Test was used to assess fluency of speech. Performance was quantified by: 1) number of words correctly listed in the test’s phonemic condition (vf_1), 2) number of words correctly listed in the test’s semantic condition (vf_2), 3) number of words correctly listed in the test’s switching condition (vf_3; Hatlestad-Hall et al., 2022). In our secondary analysis we quantified verbal fluency performance by calculating the average of vf_1, vf_2, and vf_3.

### Constructs of Cognitive Domains

Composite scores derived from the neuropsychological test battery were calculated for cognitive domains labelled as Executive Function, Psychomotor Speed, and Working Memory. A composite score representing Executive Function by averaging the time to complete the interference and switching conditions of the D-KEFS CWIT, the letter, number, and switching conditions of the D-KEFS Trail Making Test, and the correct number of words listed during the phonemic, semantic, and switching condition of the D-KEFS Verbal Fluency Test. To ensure congruency in the composite score, the timed scores were reversed scored. A composite score to represent Psychomotor Speed by averaging the time to complete the colour only and reading only conditions of the D-KEFS CWIT. A composite score to represent Working Memory by averaging the achieved score in the test’s backwards, forwards, and the sequencing condition of the WAIS-IV Digit Span Test, and the number of items correctly recalled after all five learning trials, and 30 minutes after the final learning trial of the Rey Auditory Verbal Learning Test.

### Experimental Design and Statistical Analysis

A statistical approach to reduce the complexity of the EEG data was performed using Python (v3.11.5) following the method proposed by Euler et al. (2024). Firstly, the average exponent and offset values across the scalp were plotted topographically using MNE (v1.5.1), and correlation matrices were performed to identify regions of electrodes that exhibited similar aperiodic activity (numpy v1.26.3). Following this, principal components analysis (PCA) was performed on the exponent and offset data independently to identify dominant components within the aperiodic EEG data. Lastly, K-Means clustering was performed to identify electrode clusters that exhibited distinct exponent and offset activity (skLearn v1.3.2). From this, regions of interests were constructed.

Bivariate Pearson correlations were performed to identify how the demographic, EEG, and cognitive variables associated with each other. As age has significant associations with aperiodic activity and cognitive function, partial Pearson correlations for the single neuropsychological tests and the cognitive domain constructs were also performed accounting for the age variance in the cohort (pingouin v0.5.3). Following this, we fit the data using iteratively reweighted least-squares robust regression models (statsmodels v0.14.0; Holland & Welsch, 1977) to mitigate the effects of heteroscedasticity of the data. A base formula was constructed that accounted for the impact of age:

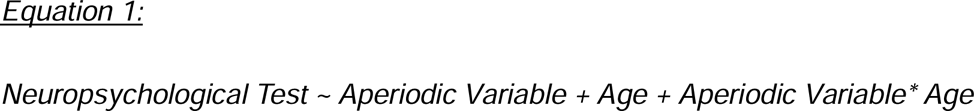

Models were constructed for each neuropsychological test and cognitive domain with separate models for the aperiodic exponent and offset. We used the “interactions” package (v1.1.5) in R (v4.3.1) to calculate simple slopes and Johnson-Neyman intervals to determine at which age aperiodic activity interactions were associated with performance of each cognitive test (-1 SD = 23.5 years, 0 SD = 37.5 years, +1 SD = 51.5 years). All p values for all tests were corrected for multiple comparisons using the Benjamini and Hochberg method (*_BH_*; Benjamini & Hochberg, 1995).

## RESULTS

### EEG Data Reduction

Topographic plots of the mean exponent and offset values were plotted to distinguish the pattern of aperiodic activity across the scalp (Figure 2A). Correlation matrices were then constructed to identify groups of electrodes that had exhibited similar relationships of aperiodic activity (Figure 2B). A qualitative inspection of correlation values across electrodes suggests a correlation of exponent values between frontal electrodes (average r = .58) and parietal electrodes (average r = .69), and a correlation of offset values between frontal electrodes (average r = .69) and parietal electrodes (average r = .67). Prior to identifying distinct cluster groups of electrodes, we constructed scalp-wide averages of the exponent and offset to represent general cognitive performance, and to see how this compares to electrode cluster regions of interest.

**Figure 2.**
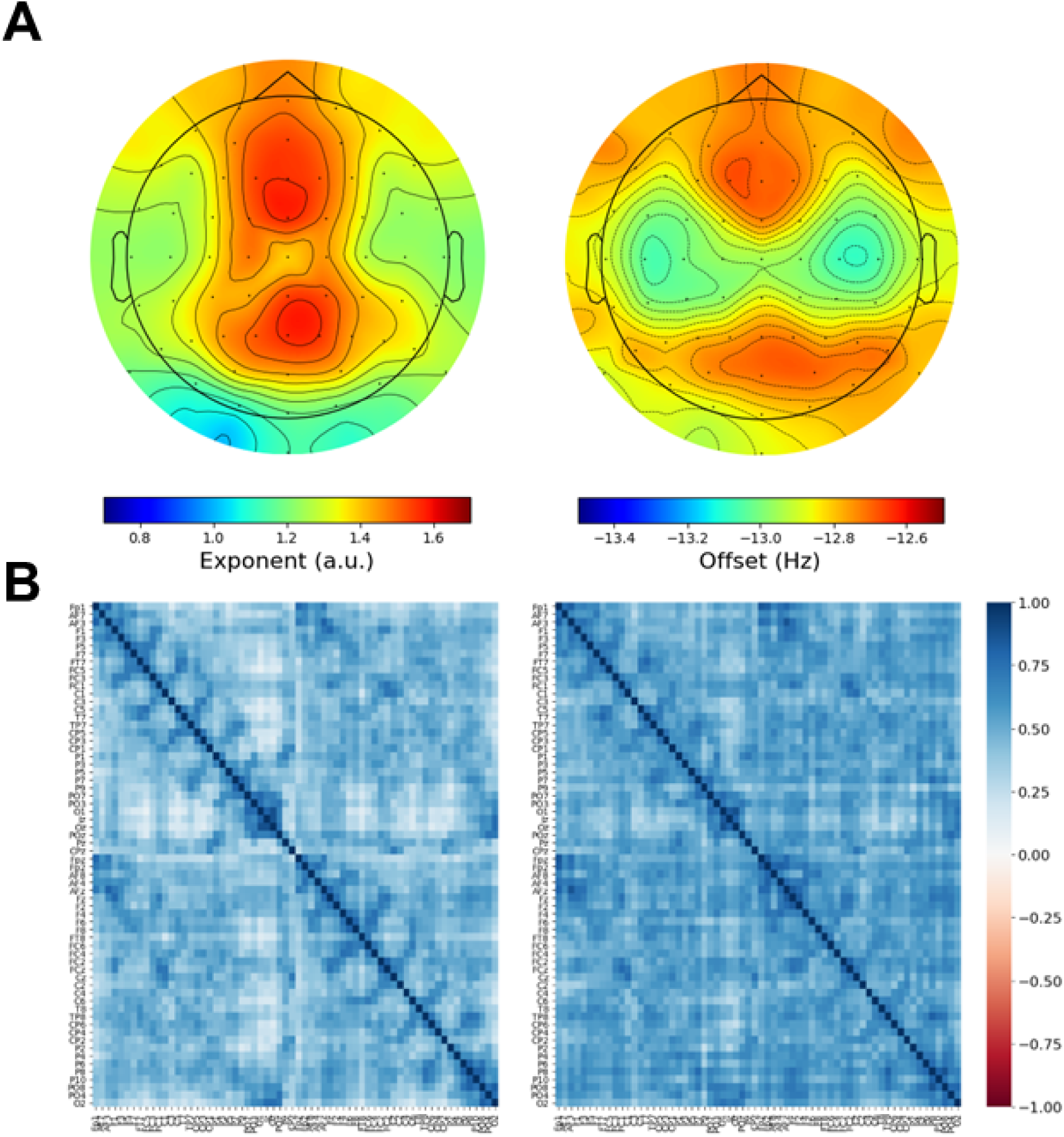
Group average topographies of the exponent and offset during eyes-closed resting EEG (A) and electrode correlation matrices for the exponent and offset (B).

PCA and K-Means clustering were then performed on the exponent (Figure 3) and offset (Figure 4) data. The PCA identified a dominant component on which electrodes loaded consistently, which accounted for 57% and 69% of the variance for the exponent and offset, respectively. The second largest component accounted for 14% and 7% of variance for the exponent and offset, respectively. This is consistent with previous PCA assessments of aperiodic activity (Euler et al., 2024). Lastly, we performed K-Means clustering to identify if there are any electrode clusters with distinct aperiodic activity. It was determined that 4 and 3 cluster centroids would be suitable for grouping the exponent (Figure 3B) and offset (Figure 4B) data in the PCA dimension space, respectively. For the exponent, frontoparietal, frontotemporal and centrotemporal, occipital, and midline clusters were identified (Figure 3C and 3D). For the offset, frontoparietal, central, and occipital clusters were identified (Figure 4C and 4D). Electrodes contained within each cluster can be found in Table 2.

**Figure 3.**
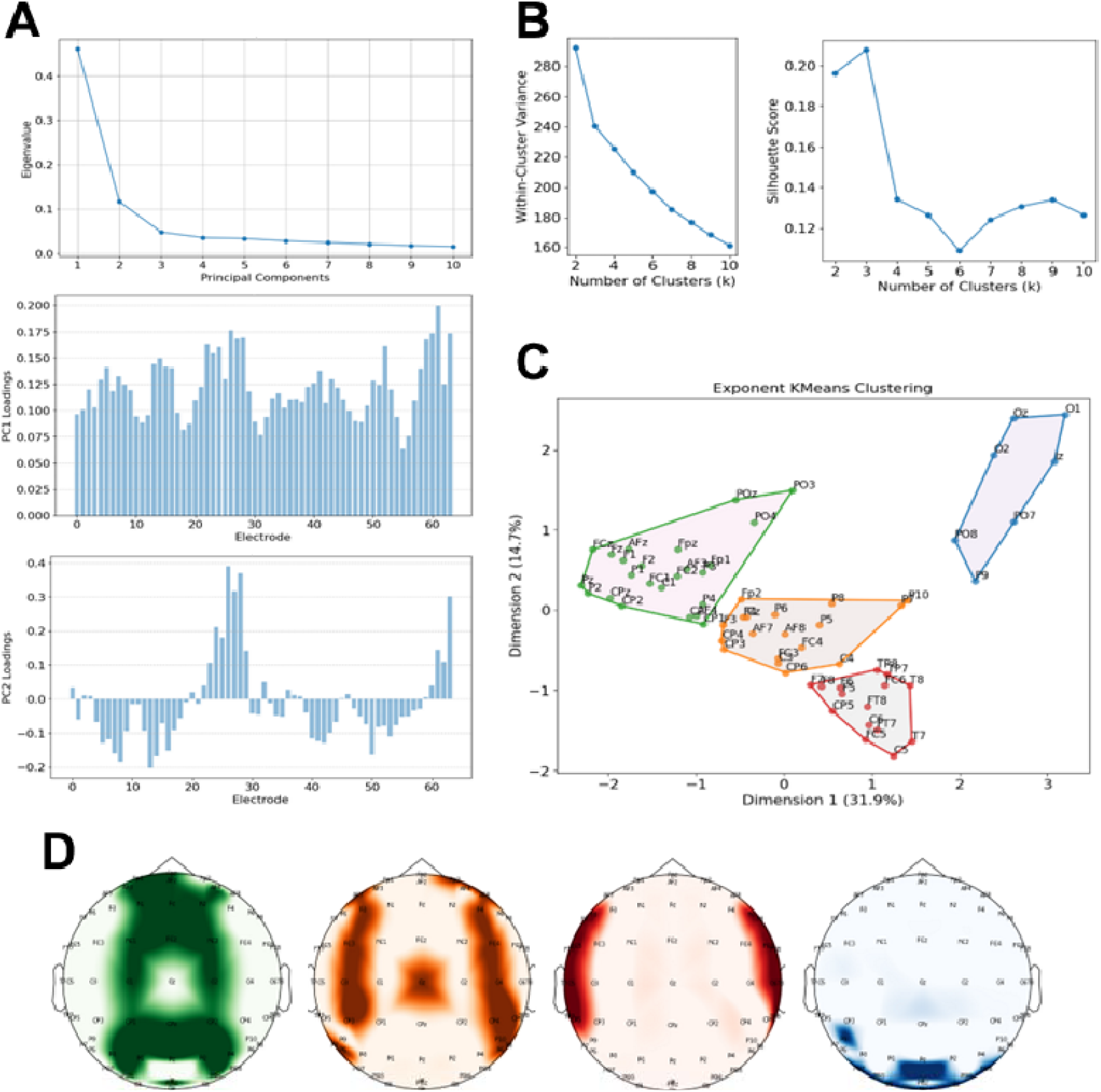
Data reduction protocol for exponent data. First, principal components analysis (PCA) was performed to identify sources of variability in the data (A). The number of clusters used in K-Means clustering was then determined by visualising the elbow plot and silhouette score of the exponent data. Four clusters were deemed suitable for clustering (B). Clustering of the exponent data was performed in the PCA dimensional space (C). Midline (green), frontoparietal (orange), frontotemporal and centrotemporal (red), and occipital (blue) electrode clusters were identified (D).

**Figure 4.**
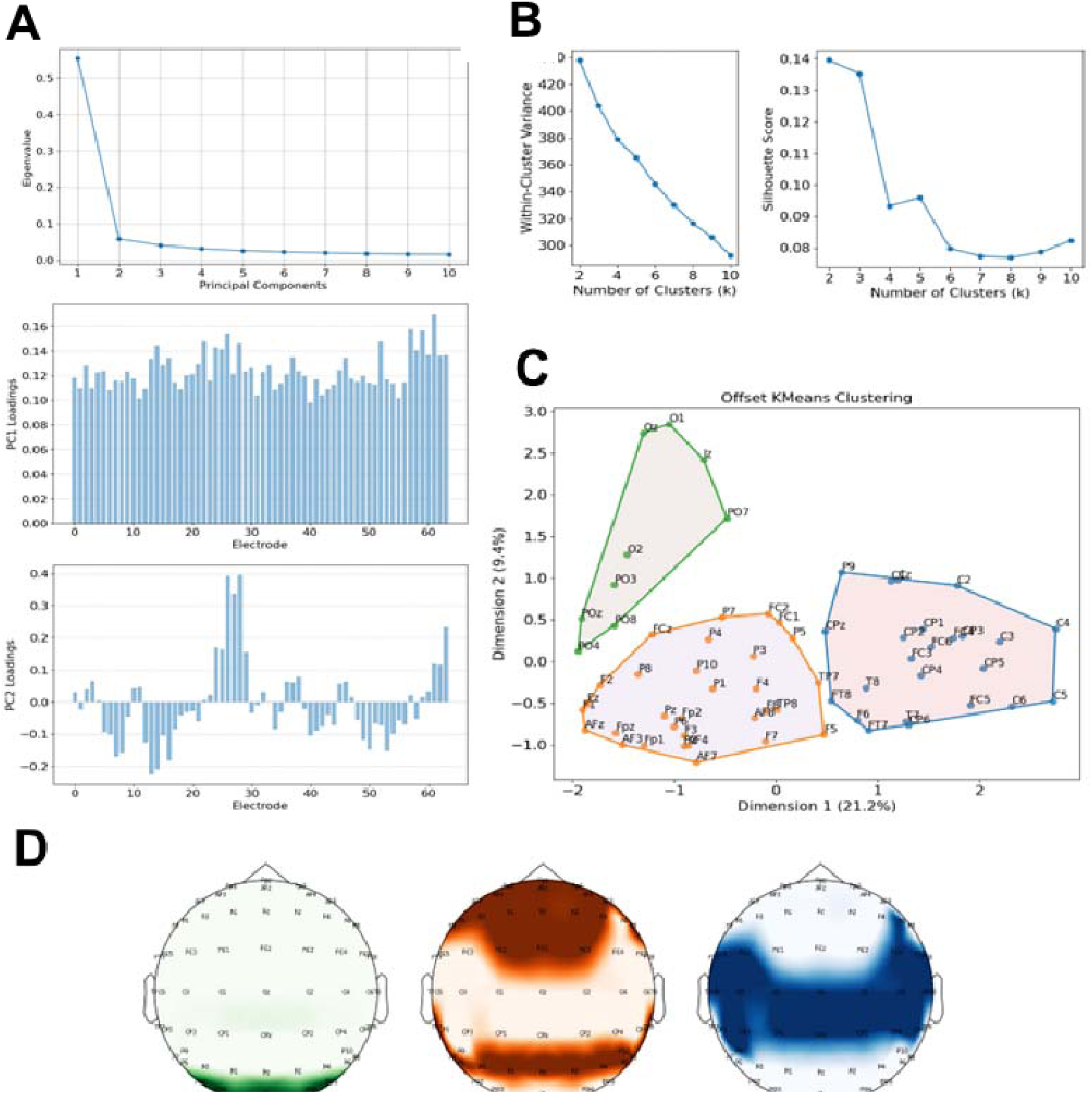
Data reduction protocol for offset data. First, principal components analysis (PCA) was performed to identify sources of variability in the data (A). The number of clusters used in K-Means clustering was then determined by visualising the elbow plot and silhouette score of the offset data. Three clusters were deemed suitable for clustering (B). Clustering of the offset data was performed in the PCA dimensional space (C). Occipital (green), frontoparietal (orange), and central (blue) electrode clusters were identified (D).

**Table 2:** Electrodes contained within each cluster.

| **Electrodes** |  |
| --- | --- |
| ***Exponent clusters*** |  |
| Frontoparietal | Fp1, AF7, AF3, F1, F3, F5, F7, FC1, TP7, P1, P3, P5, P7, Pz, Fpz, Fp2, AF8, AF4, AFz, Fz, F2, F4, F8, FC2, FCz, TP8, P2, P4, P6, P8, P10 |
| Occipital | PO7, PO3, O1, Iz, Oz, POz, PO8, PO4, O2 |
| Midline | Fp1, AF3, F1, FC1, C1, CP1, P1, P3, PO3, POz, Pz, CPz, Fpz, AF4, AFz, Fz, F2, FC2, FCz, C2, CP2, P2, P4, PO4 |
| Fronto/Centrotemporal | FT7, FC5, FC3, C1, C3, C5, T7, CP5, CP3, CP1, P9, CPz, F6, FT8, FC6, FC4, Cz, C2, C4, C6, T8, CP6, CP4, CP2 |
| ***Offset clusters*** |  |
| Frontoparietal | Fp1, AF7, AF3, F1, F3, F5, F7, FC1, TP7, P1, P3, P5, P7, Pz, Fpz, Fp2, AF8, AF4, AFz, Fz, F2, F4, F8, FC2, FCz, TP8, P2, P4, P6, P8, P10 |
| Occipital | PO7, PO3, O1, Iz, Oz, POz, PO8, PO4, O2 |
| Central | FT7, FC5, FC3, C1, C3, C5, T7, CP5, CP3, CP1, P9, CPz, F6, FT8, FC6, FC4, Cz, C2, C4, C6, T8, CP6, CP4, CP2 |

### Correlations

Results of the zero-order correlations are available in the top right quadrant of Table 3. Results of the partial correlations accounting for age are available in the bottom left quadrant of Table 3 and significant correlations are summarised below.

**Table 3:**
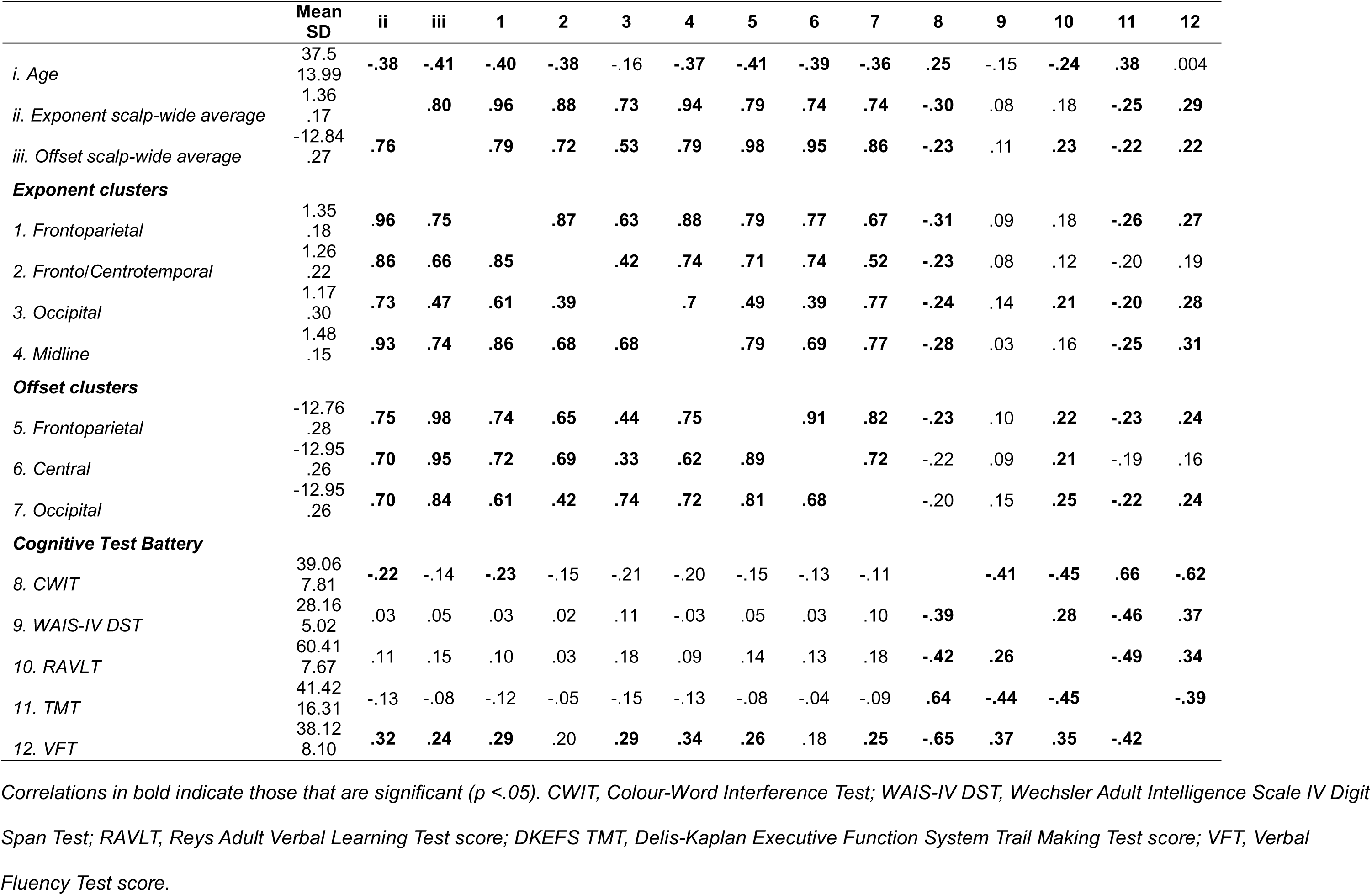
Descriptive statistics, bivariate correlation (upper right quadrant), and partial correlations controlling for age (lower left quadrant).

|  | Mean<br>SD | ii | iii | 1 | 2 | 3 | 4 | 5 | 6 | 7 | 8 | 9 | 10 | 11 | 12 |
| --- | --- | --- | --- | --- | --- | --- | --- | --- | --- | --- | --- | --- | --- | --- | --- |
| *i. Age* | 37.5<br>13.99 | **-.38** | **-.41** | **-.40** | **-.38** | -.16 | **-.37** | **-.41** | **-.39** | **-.36** | **.25** | -.15 | **-.24** | **.38** | .004 |
| *ii. Exponent scalp-wide average* | 1.36<br>.17 |  | **.80** | **.96** | **.88** | **.73** | **.94** | **.79** | **.74** | **.74** | **-.30** | .08 | .18 | **-.25** | **.29** |
| *iii. Offset scalp-wide average* | -12.84<br>.27 | **.76** |  | **.79** | **.72** | **.53** | **.79** | **.98** | **.95** | **.86** | **-.23** | .11 | **.23** | **-.22** | **.22** |
| **Exponent clusters** |  |  |  |  |  |  |  |  |  |  |  |  |  |  |  |
| *1. Frontoparietal* | 1.35<br>.18 | **.96** | **.75** |  | **.87** | **.63** | **.88** | **.79** | **.77** | **.67** | **-.31** | .09 | .18 | **-.26** | **.27** |
| *2. Fronto/Centrotemporal* | 1.26<br>.22 | **.86** | **.66** | **.85** |  | **.42** | **.74** | **.71** | **.74** | **.52** | **-.23** | .08 | .12 | -.20 | .19 |
| *3. Occipital* | 1.17<br>.30 | **.73** | **.47** | **.61** | **.39** |  | **.7** | **.49** | **.39** | **.77** | **-.24** | .14 | **.21** | **-.20** | **.28** |
| *4. Midline* | 1.48<br>.15 | **.93** | **.74** | **.86** | **.68** | **.68** |  | **.79** | **.69** | **.77** | **-.28** | .03 | .16 | **-.25** | **.31** |
| **Offset clusters** |  |  |  |  |  |  |  |  |  |  |  |  |  |  |  |
| *5. Frontoparietal* | -12.76<br>.28 | **.75** | **.98** | **.74** | **.65** | **.44** | **.75** |  | **.91** | **.82** | **-.23** | .10 | **.22** | **-.23** | **.24** |
| *6. Central* | -12.95<br>.26 | **.70** | **.95** | **.72** | **.69** | **.33** | **.62** | **.89** |  | **.72** | -.22 | .09 | **.21** | -.19 | .16 |
| *7. Occipital* | -12.95<br>.26 | **.70** | **.84** | **.61** | **.42** | **.74** | **.72** | **.81** | **.68** |  | -.20 | .15 | **.25** | **-.22** | **.24** |
| **Cognitive Test Battery** |  |  |  |  |  |  |  |  |  |  |  |  |  |  |  |
| *8. CWIT* | 39.06<br>7.81 | **-.22** | -.14 | **-.23** | -.15 | -.21 | -.20 | -.15 | -.13 | -.11 |  | **-.41** | **-.45** | **.66** | **-.62** |
| *9. WAIS-IV DST* | 28.16<br>5.02 | .03 | .05 | .03 | .02 | .11 | -.03 | .05 | .03 | .10 | **-.39** |  | **.28** | **-.46** | **.37** |
| *10. RAVLT* | 60.41<br>7.67 | .11 | .15 | .10 | .03 | .18 | .09 | .14 | .13 | .18 | **-.42** | **.26** |  | **-.49** | **.34** |
| *11. TMT* | 41.42<br>16.31 | -.13 | -.08 | -.12 | -.05 | -.15 | -.13 | -.08 | -.04 | -.09 | **.64** | **-.44** | **-.45** |  | **-.39** |
| *12. VFT* | 38.12<br>8.10 | **.32** | **.24** | **.29** | .20 | **.29** | **.34** | **.26** | .18 | **.25** | **-.65** | **.37** | **.35** | **-.42** |  |

*Correlations in bold indicate those that are significant ( $p < .05$ ). CWIT, Colour-Word Interference Test; WAIS-IV DST, Wechsler Adult Intelligence Scale IV Digit Span Test; RAVLT, Reys Adult Verbal Learning Test score; DKEFS TMT, Delis-Kaplan Executive Function System Trail Making Test score; VFT, Verbal Fluency Test score.*

Age was negatively correlated with the average exponent (*r* = -.38, *p_BH_* <.001) and the average offset (*r* = -.41, *p_BH_* <.001), and at frontoparietal (*r* = -.40, *p_BH_* <.001), frontotemporal and centrotemporal (*r* = -.38, *p_BH_* <.001), and midline clusters (*r* = -.37, *p_BH_* <.001) for the exponent, and at frontoparietal (*r* = -.41, *p_BH_* <.001), central (*r* = -.39, *p_BH_* <.001), and occipital (*r* = -.36, *p_BH_* <.001) clusters for the offset. That is, the exponent and the offset reduced with age. Age was associated with longer durations to perform the Colour-Word Interference Test (*r* = .25, *p_BH_* = .012) and the Trail Making Test (*r* = .38, *p_BH_* = <.001), and lower scores on the Rey Adult Verbal Learning Test (*r* = .24, *p_BH_*= <.001).

When accounting for age, the exponent at each electrode cluster correlated with each other (*r* = .39 – .86, *p_BH_* < .001), as well as with the scalp-wide average exponent (*r* = .73 – .96, *p_BH_* < .001). The scalp-wide average exponent (*r* = -.22, CI = -.41 – -.05, *p_BH_* = .034) and frontoparietal exponent (*r* = -.23, CI = -.41 – -.05, *p_BH_* = .026) negatively correlated with the Colour-Word Interference Test. The scalp-wide average exponent (*r* = .32, CI = .14 – .48, *p_BH_* = .001), frontoparietal exponent (*r* = .29, CI = .11 – .45, *p_BH_* = .004), occipital exponent (*r* = .29, CI = .1 – .45, *p_BH_* = .005), and midline exponent (*r* = .34, CI = .16 – .49, *p_BH_* < .001), were all positively correlated with Verbal Fluency Test performance. That is, greater exponents were associated with faster time during the Colour-Word Interference Test and greater score during the Verbal Fluency Test.

The offset at each electrode cluster correlated with each other (*r* = .68 - .89, *p_BH_* < .001), as well as with the scalp-wide average offset (*r* = .84 – .98, *p_BH_* < .001). The frontoparietal offset (*r* = .26, CI = .08 – .43, *p_BH_* = .012), and the occipital offset (*r* = .25, CI = .07 – .42, *p_BH_* = .015) positively correlated with Verbal Fluency Test performance. That is, higher offsets were associated with better performance during the Verbal Fluency Test.

When controlling for age, the average exponent (*r* = .22, *p*_BH_ = .03), frontoparietal exponent (r = .24, *p*_BH_ = .02) and midline exponent (r = .21, *p*_BH_ = .04) were positively correlated with Executive Function. Similarly, the average exponent (*r* = -.22, *p*_BH_ = .03), frontoparietal exponent (r = -.22, *p* = .03), and occipital exponent (r = -.26, *p*_BH_ = .009) were negatively correlated with Psychomotor Speed. Whereas Working Memory was only positively correlated with the occipital exponent (*r* = .23, *p*_BH_ = .02). That is, greater exponents were associated with better Executive Function and faster Psychomotor Speed across the scalp, whereas only associated with better Working Memory at occipital locations. There were no significant correlations between the aperiodic offset and the cognitive domain constructs Executive Function, Psychomotor Speed, nor Working Memory.

### Aperiodic Activity and Age as Predictors of Cognitive Function

There were only significant main effects for the Colour-Word Interference Test, whereas for the Verbal Fluency Test there were a main effect and an interaction effect. Results of these tests are summarised below while the results of the models for the other tests, are reported in Table 4 for the exponent models and Table 5 for the offset models.

**Table 4:**
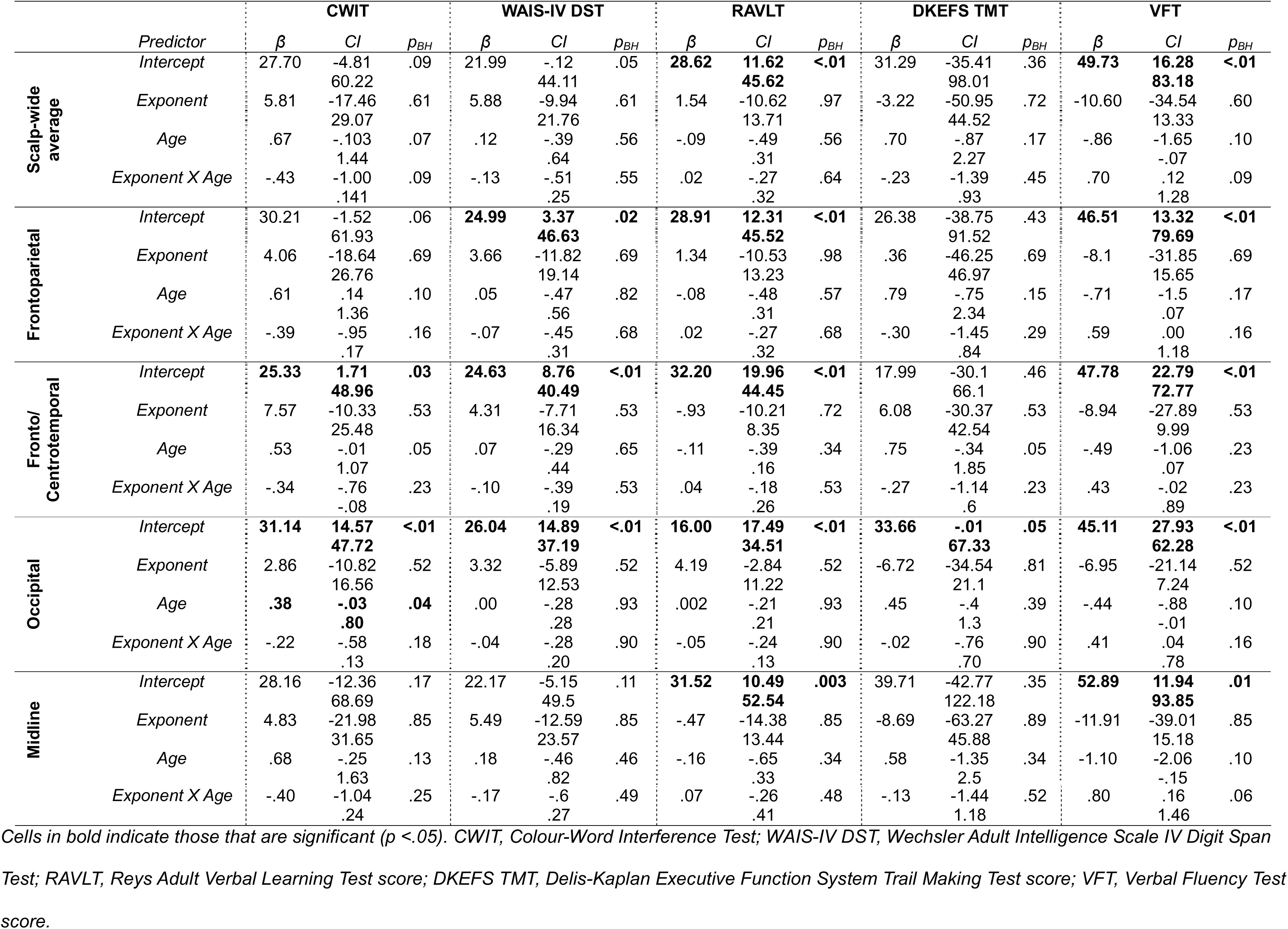
Exponent by age robust regression models for individual neuropsychological tests.

|  |  | CWIT |  |  | WAIS-IV DST |  |  | RAVLT |  |  | DKEFS TMT |  |  | VFT |  |  |
| --- | --- | --- | --- | --- | --- | --- | --- | --- | --- | --- | --- | --- | --- | --- | --- | --- |
| *Predictor* | | $\beta$ | *CI* | $p_{BH}$ | $\beta$ | *CI* | $p_{BH}$ | $\beta$ | *CI* | $p_{BH}$ | $\beta$ | *CI* | $p_{BH}$ | $\beta$ | *CI* | $p_{BH}$ |
| Scalp-wide average | *Intercept* | 27.70 | -4.81<br>60.22 | .09 | 21.99 | -.12<br>44.11 | .05 | **28.62** | **11.62**<br>**45.62** | **<.01** | 31.29 | -35.41<br>98.01 | .36 | **49.73** | **16.28**<br>**83.18** | **<.01** |
|  | *Exponent* | 5.81 | -17.46<br>29.07 | .61 | 5.88 | -9.94<br>21.76 | .61 | 1.54 | -10.62<br>13.71 | .97 | -3.22 | -50.95<br>44.52 | .72 | -10.60 | -34.54<br>13.33 | .60 |
|  | *Age* | .67 | -.103<br>1.44 | .07 | .12 | -.39<br>.64 | .56 | -.09 | -.49<br>.31 | .56 | .70 | -.87<br>2.27 | .17 | -.86 | -1.65<br>-.07 | .10 |
|  | *Exponent X Age* | -.43 | -1.00<br>.141 | .09 | -.13 | -.51<br>.25 | .55 | .02 | -.27<br>.32 | .64 | -.23 | -1.39<br>.93 | .45 | .70 | .12<br>1.28 | .09 |
| Frontoparietal | *Intercept* | 30.21 | -1.52<br>61.93 | .06 | **24.99** | **3.37**<br>**46.63** | **.02** | **28.91** | **12.31**<br>**45.52** | **<.01** | 26.38 | -38.75<br>91.52 | .43 | **46.51** | **13.32**<br>**79.69** | **<.01** |
|  | *Exponent* | 4.06 | -18.64<br>26.76 | .69 | 3.66 | -11.82<br>19.14 | .69 | 1.34 | -10.53<br>13.23 | .98 | .36 | -46.25<br>46.97 | .69 | -8.1 | -31.85<br>15.65 | .69 |
|  | *Age* | .61 | .14<br>1.36 | .10 | .05 | -.47<br>.56 | .82 | -.08 | -.48<br>.31 | .57 | .79 | -.75<br>2.34 | .15 | -.71 | -1.5<br>.07 | .17 |
|  | *Exponent X Age* | -.39 | -.95<br>.17 | .16 | -.07 | -.45<br>.31 | .68 | .02 | -.27<br>.32 | .68 | -.30 | -1.45<br>.84 | .29 | .59 | .00<br>1.18 | .16 |
| Fronto/ Centrotemporal | *Intercept* | **25.33** | **1.71**<br>**48.96** | **.03** | **24.63** | **8.76**<br>**40.49** | **<.01** | **32.20** | **19.96**<br>**44.45** | **<.01** | 17.99 | -30.1<br>66.1 | .46 | **47.78** | **22.79**<br>**72.77** | **<.01** |
|  | *Exponent* | 7.57 | -10.33<br>25.48 | .53 | 4.31 | -7.71<br>16.34 | .53 | -.93 | -10.21<br>8.35 | .72 | 6.08 | -30.37<br>42.54 | .53 | -8.94 | -27.89<br>9.99 | .53 |
|  | *Age* | .53 | -.01<br>1.07 | .05 | .07 | -.29<br>.44 | .65 | -.11 | -.39<br>.16 | .34 | .75 | -.34<br>1.85 | .05 | -.49 | -1.06<br>.07 | .23 |
|  | *Exponent X Age* | -.34 | -.76<br>-.08 | .23 | -.10 | -.39<br>.19 | .53 | .04 | -.18<br>.26 | .53 | -.27 | -1.14<br>.6 | .23 | .43 | -.02<br>.89 | .23 |
| Occipital | *Intercept* | **31.14** | **14.57**<br>**47.72** | **<.01** | **26.04** | **14.89**<br>**37.19** | **<.01** | **16.00** | **17.49**<br>**34.51** | **<.01** | **33.66** | **-.01**<br>**67.33** | **.05** | **45.11** | **27.93**<br>**62.28** | **<.01** |
|  | *Exponent* | 2.86 | -10.82<br>16.56 | .52 | 3.32 | -5.89<br>12.53 | .52 | 4.19 | -2.84<br>11.22 | .52 | -6.72 | -34.54<br>21.1 | .81 | -6.95 | -21.14<br>7.24 | .52 |
|  | *Age* | **.38** | **-.03**<br>**.80** | **.04** | .00 | -.28<br>.28 | .93 | .002 | -.21<br>.21 | .93 | .45 | -.4<br>1.3 | .39 | -.44 | -.88<br>-.01 | .10 |
|  | *Exponent X Age* | -.22 | -.58<br>.13 | .18 | -.04 | -.28<br>.20 | .90 | -.05 | -.24<br>.13 | .90 | -.02 | -.76<br>.70 | .90 | .41 | .04<br>.78 | .16 |
| Midline | *Intercept* | 28.16 | -12.36<br>68.69 | .17 | 22.17 | -5.15<br>49.5 | .11 | **31.52** | **10.49**<br>**52.54** | **.003** | 39.71 | -42.77<br>122.18 | .35 | **52.89** | **11.94**<br>**93.85** | **.01** |
|  | *Exponent* | 4.83 | -21.98<br>31.65 | .85 | 5.49 | -12.59<br>23.57 | .85 | -.47 | -14.38<br>13.44 | .85 | -8.69 | -63.27<br>45.88 | .89 | -11.91 | -39.01<br>15.18 | .85 |
|  | *Age* | .68 | -.25<br>1.63 | .13 | .18 | -.46<br>.82 | .46 | -.16 | -.65<br>.33 | .34 | .58 | -1.35<br>2.5 | .34 | -1.10 | -2.06<br>-.15 | .10 |

|  |  |  |  |  |  |  |  |  |  |  |  |  |  |  |  |
| --- | --- | --- | --- | --- | --- | --- | --- | --- | --- | --- | --- | --- | --- | --- | --- |
| *Exponent X Age* | *-.40* | *-1.04* | *.25* | *-.17* | *-.6* | *.49* | *.07* | *-.26* | *.48* | *-.13* | *-1.44* | *.52* | *.80* | *.16* | *.06* |
|  |  | *.24* |  |  | *.27* |  |  | *.41* |  |  | *1.18* |  |  | *1.46* |  |

Cells in bold indicate those that are significant ( $p < .05$ ). *CWIT*, Colour-Word Interference Test; *WAIS-IV DST*, Wechsler Adult Intelligence Scale IV Digit Span Test; *RAVLT*, Reys Adult Verbal Learning Test score; *DKEFS TMT*, Delis-Kaplan Executive Function System Trail Making Test score; *VFT*, Verbal Fluency Test score.

**Table 5:** Offset by age robust regression models for individual neuro psychologic al tests.

|  | *Predictor* | CWIT |  |  | WAIS-IV DST |  |  | RAVLT |  |  | DKEFS TMT |  |  | VFT |  |  |
| --- | --- | --- | --- | --- | --- | --- | --- | --- | --- | --- | --- | --- | --- | --- | --- | --- |
| | | $\beta$ | *CI* | $p_{BH}$ | $\beta$ | *CI* | $p_{BH}$ | $\beta$ | *CI* | $p_{BH}$ | $\beta$ | *CI* | $p_{BH}$ | $\beta$ | *CI* | $p_{BH}$ |
| Scalp-wide average | *Intercept* | 52.61 | -123.64<br>228.86 | .56 | 36.89 | -80.49<br>154.28 | .54 | 39.99 | -49.49<br>129.47 | .38 | 126.25 | -228.01<br>480.52 | .49 | -95.91 | -275.26<br>83.44 | .29 |
|  | *Offset* | 1.36 | -12.38<br>15.11 | .96 | .55 | -8.61<br>9.71 | .96 | .74 | -6.24<br>7.72 | .96 | 7.77 | -19.86<br>35.41 | .68 | -10.22 | -24.21<br>3.77 | .67 |
|  | *Age* | -1.89 | -6.22<br>2.42 | .82 | .14 | -2.73<br>3.03 | .96 | .50 | -1.68<br>2.70 | .78 | -3.88 | -12.57<br>4.81 | .30 | **6.38** | **1.98**<br>**10.78** | **.02** |
|  | *Offset X Age* | -.15 | -.49<br>.18 | .76 | .01 | -.21<br>.24 | .94 | .04 | -.13<br>.21 | .73 | -.33 | -1.01<br>.34 | .24 | **.48** | **.15**<br>**.83** | **.02** |
| Frontoparietal | *Intercept* | 28.56 | -141.07<br>198.19 | .74 | 39.68 | -73.18<br>152.53 | .49 | 44.11 | -42.06<br>130.28 | .32 | 97.78 | -242.86<br>438.43 | .57 | -68.51 | -240.86<br>103.83 | .44 |
|  | *Offset* | -.51 | -13.82<br>12.79 | .81 | .76 | -8.09<br>9.82 | .88 | 1.06 | -5.69<br>7.82 | .88 | 5.58 | -21.14<br>32.31 | .79 | -8.13 | -21.64<br>5.4 | .79 |
|  | *Age* | -1.18 | -5.26<br>2.89 | .93 | .01 | -2.71<br>2.72 | .93 | .29 | -1.77<br>2.37 | .93 | -3.1 | -11.28<br>5.09 | .29 | **5.59** | **1.44**<br>**9.73** | **.04** |
|  | *Offset X Age* | -.10 | -.42<br>.22 | .95 | .01 | -.21<br>.22 | .95 | .02 | -.13<br>.18 | .94 | -.27 | -.91<br>.37 | .23 | **.43** | **.11**<br>**.75** | **.04** |
| Central | *Intercept* | 79.65 | -101.38<br>260.69 | .39 | 27.91 | -92.81<br>148.64 | .65 | 16.89 | -75.08<br>108.87 | .72 | 172.55 | -191.76<br>536.49 | .35 | -119.82 | -307.58<br>67.94 | .21 |
|  | *Offset* | 3.45 | -10.55<br>17.44 | .97 | -.15 | -9.45<br>9.17 | .97 | -1.05 | -8.16<br>6.05 | .97 | 11.32 | -16.84<br>39.49 | .47 | -12.04 | -26.56<br>2.48 | .47 |
|  | *Age* | -2.57 | -7.17<br>2.01 | .61 | .23 | -2.83<br>3.29 | .91 | 1.04 | -1.28<br>3.38 | .45 | -4.64 | -13.88<br>4.6 | .27 | **6.49** | **1.74**<br>**11.26** | **.04** |
|  | *Offset X Age* | -.21 | -.56<br>.14 | .56 | .02 | -.21<br>.26 | .88 | .08 | -.09<br>.26 | .41 | -.39 | -1.1<br>.32 | .22 | **.5** | **.13**<br>**.86** | **.04** |
| Occipital | *Intercept* | 70.96 | -76.25<br>218.17 | .34 | 40.15 | -57.26<br>137.56 | .42 | 60.31 | -13.97<br>134.60 | .08 | 86.76 | -208.14<br>381.68 | .56 | -85.91 | -234.46<br>62.64 | .26 |
|  | *Offset* | 2.83 | -8.69<br>14.36 | .83 | .82 | -6.80<br>8.45 | .83 | 2.33 | -3.48<br>8.14 | .83 | 4.71 | -18.37<br>27.8 | .83 | -9.51 | -21.14<br>2.12 | .47 |
|  | *Age* | -1.83 | -5.47<br>1.81 | .73 | .25 | -2.16<br>2.66 | .82 | -.09 | -1.93<br>1.74 | .77 | -2.82 | -10.13<br>4.47 | .72 | **5.58** | **1.91**<br>**9.26** | **.01** |
|  | *Offset X Age* | -0.15 | -.43<br>.13 | .64 | .02 | -.17<br>.21 | .77 | -.003 | -.14<br>.14 | .77 | -.25 | -.82<br>.32 | .59 | **.43** | **.14**<br>**.72** | **.01** |

*Cells in bold indicate those that are significant ( $p < .05$ ). CWIT, Colour-Word Interference Test; WAIS-IV DST, Wechsler Adult Intelligence Scale IV Digit Span Test; RAVLT, Reys Adult Verbal Learning Test score; DKEFS TMT, Delis-Kaplan Executive Function System Trail Making Test score; VFT, Verbal Fluency Test score.*

#### Colour-Word Interference Test

In the exponent models, there was a main effect of age for the occipital exponent (β = .38, CI = -.03 – .80, *p_BH_* = .049). There were no other main effects or interactions for the scalp-wide average exponent nor the exponent clusters.

#### Verbal Fluency Test

In the offset models, there was a main effect of age for the scalp-wide average (β = 6.38, CI = 1.98 – 10.78, *p_BH_*= .022), and at frontoparietal (β = 5.59, CI = 1.44 – 9.73, *p_BH_* = .041), central (β = 6.49, CI = 1.74 – 11.26, *p_BH_* = .044), and occipital (β = 5.58, CI = 1.91 – 9.26, *p_BH_* = .017) clusters, which were moderated by age interactions for the scalp-wide average (β = .48, CI = .15 – .83, *p_BH_* = .024), and for the frontoparietal (β = .43, CI = .11 – .75, *p_BH_* = .045), central (β = .5, CI = .13 – .86, *p_BH_* = .046), and occipital (β = .43, CI = .14 – .72, *p_BH_* = .018) clusters. Johnson-Neyman plots revealed that associations were significant from 33 – 37 years of age. Simple slopes revealed that the scalp-wide association was not significant at -1 SD of age (β = 1.29, *p* = .740) while significant at +1 SD of age (β = 15.00, *p* < .001; Figure 5A and 5B). The frontoparietal cluster association was not significant at -1 SD of age (β = .39, *p* = .920) while significant at +1 SD of age (β = 13.49, *p* < .001; Figure 5C and 5D). The central cluster association was not significant at -1 SD of age (β = 1.65, *p* = .665) while significant at +1 SD of age (β = 13.93, *p* < .001; Figure 5E and 5F). The occipital cluster association was not significant at -1 SD of age (β = .60, *p* = .850) while significant at +1 SD of age (β = 12.64, *p* < .001; Figure 5G and 5H).

**Figure 5.**
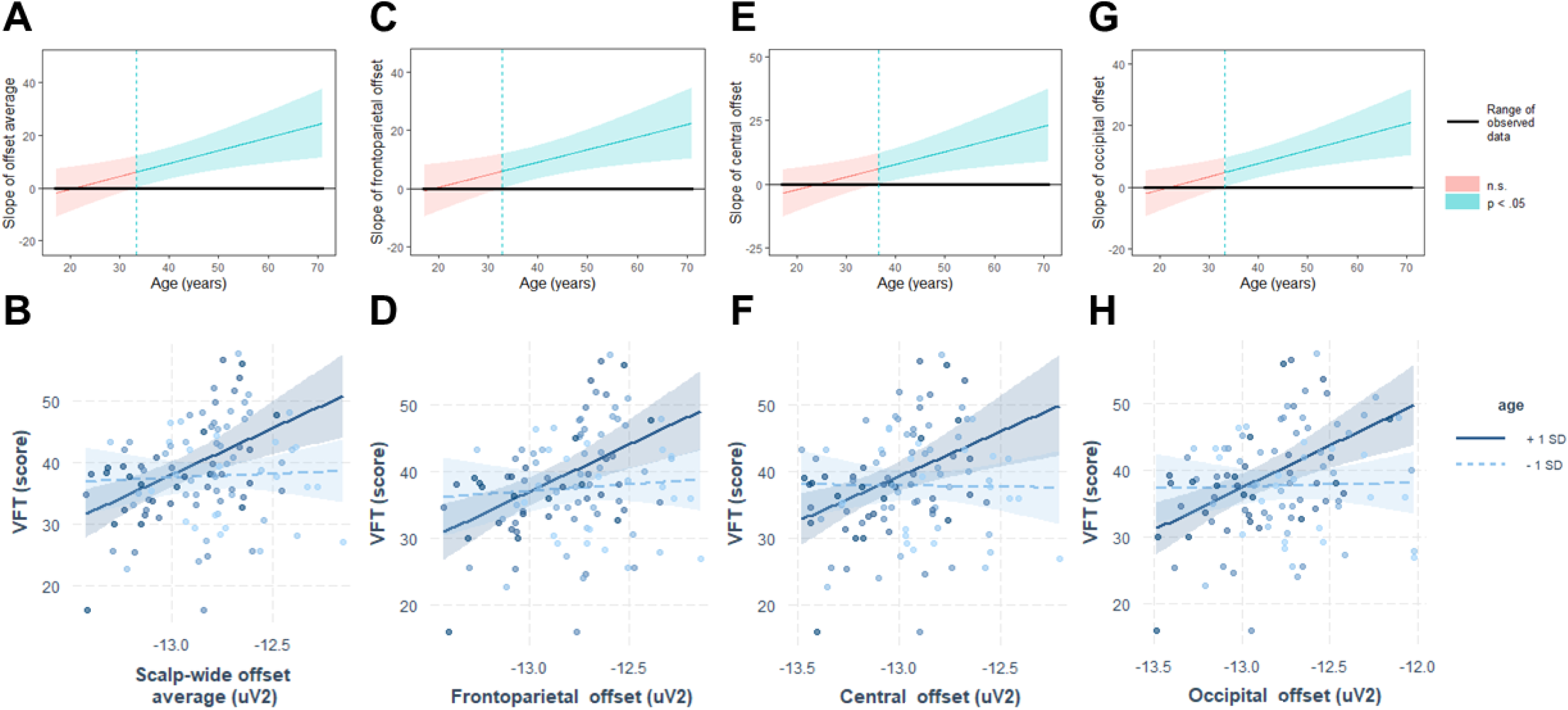
Johnson-Neyman plots (top row) and interaction plots (bottom row) for Verbal Fluency Test (VFT). Reductions in the scalp-wide average (A, B), frontoparietal (C, D), central (E, F), and occipital offset were associated with worse performance during the VFT for adults over the age of 33-37 years. Greater score indicates improving performance during the VFT. -1 SD age = 23.5 years, +1 SD age = 51.5 years. Error bars represent 95% confidence intervals.

#### Cognitive Domain constructs

In the exponent models (Table 6), there was main effect of age for the fronto/centrotemporal cluster and Executive Function (β = - .65, CI = -1.35 – .04, *p_BH_* = .02). There were no other main effects or interaction effects for the scalp-wide average exponent nor the exponent clusters. In the offset models (Table 7), there were no main effects of interaction effects for the scalp-wide average offset nor the offset clusters.

**Table 6.**
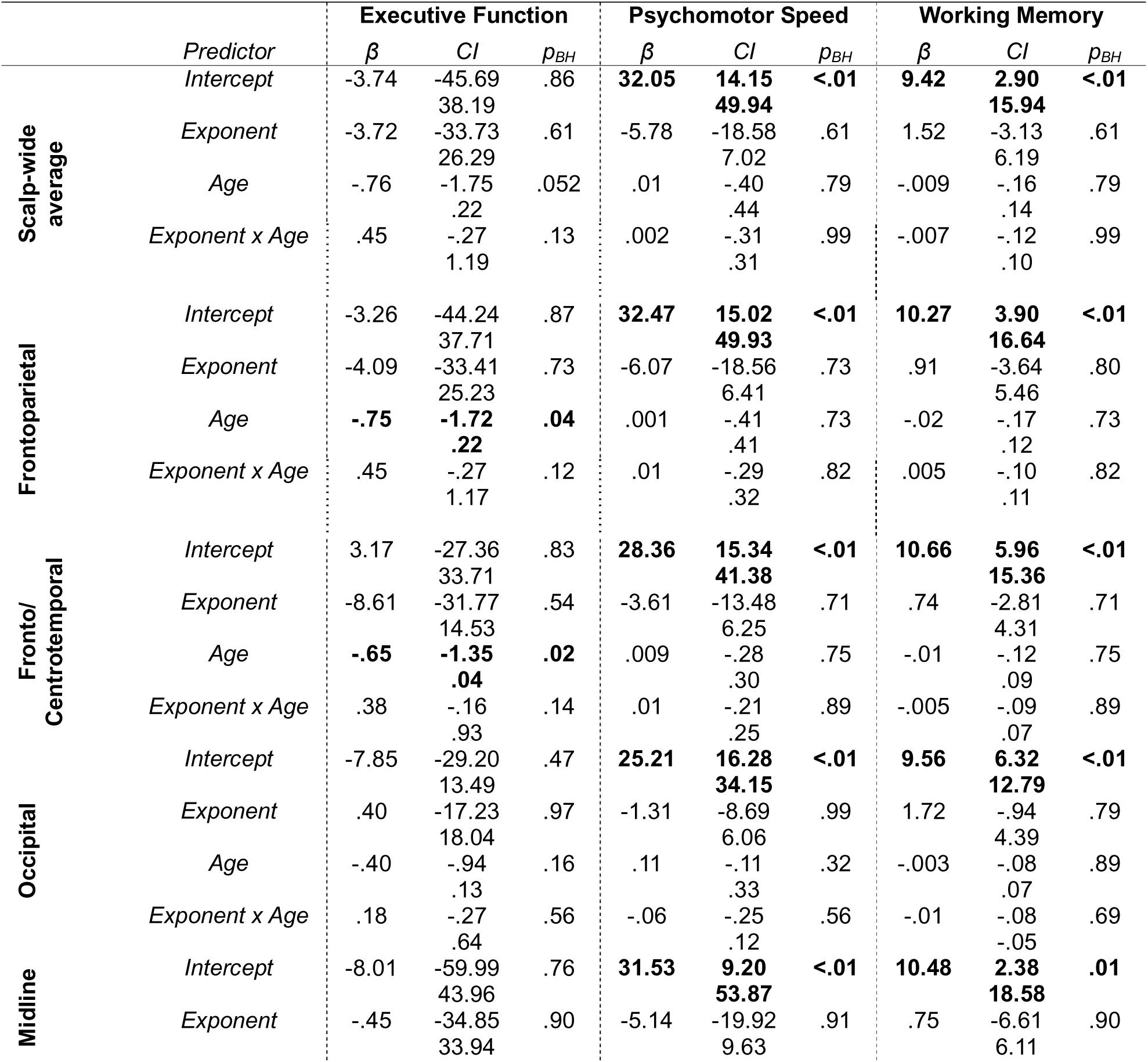
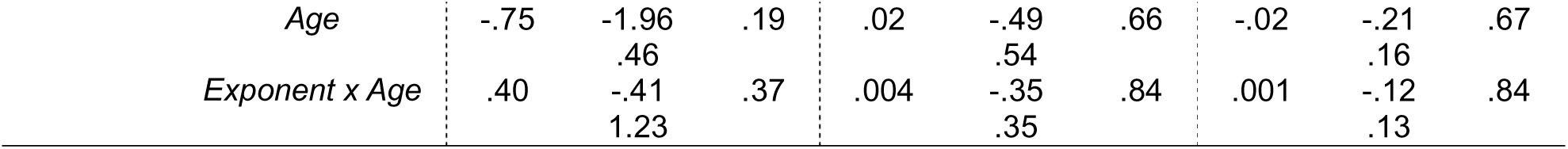
Exponent by age robust regression models for cognitive domains.

|  | *Predictor* | **Executive Function** |  |  | **Psychomotor Speed** |  |  | **Working Memory** |  |  |
| --- | --- | --- | --- | --- | --- | --- | --- | --- | --- | --- |
| | | $\beta$ | *CI* | $p_{BH}$ | $\beta$ | *CI* | $p_{BH}$ | $\beta$ | *CI* | $p_{BH}$ |
| Scalp-wide average | *Intercept* | -3.74 | -45.69<br>38.19 | .86 | **32.05** | **14.15**<br>**49.94** | **<.01** | **9.42** | **2.90**<br>**15.94** | **<.01** |
|  | *Exponent* | -3.72 | -33.73<br>26.29 | .61 | -5.78 | -18.58<br>7.02 | .61 | 1.52 | -3.13<br>6.19 | .61 |
|  | *Age* | -.76 | -1.75<br>.22 | .052 | .01 | -.40<br>.44 | .79 | -.009 | -.16<br>.14 | .79 |
|  | *Exponent x Age* | .45 | -.27<br>1.19 | .13 | .002 | -.31<br>.31 | .99 | -.007 | -.12<br>.10 | .99 |
| Frontoparietal | *Intercept* | -3.26 | -44.24<br>37.71 | .87 | **32.47** | **15.02**<br>**49.93** | **<.01** | **10.27** | **3.90**<br>**16.64** | **<.01** |
|  | *Exponent* | -4.09 | -33.41<br>25.23 | .73 | -6.07 | -18.56<br>6.41 | .73 | .91 | -3.64<br>5.46 | .80 |
|  | *Age* | **-.75** | **-1.72**<br>**.22** | **.04** | .001 | -.41<br>.41 | .73 | -.02 | -.17<br>.12 | .73 |
|  | *Exponent x Age* | .45 | -.27<br>1.17 | .12 | .01 | -.29<br>.32 | .82 | .005 | -.10<br>.11 | .82 |
| Fronto/<br>Centrotemporal | *Intercept* | 3.17 | -27.36<br>33.71 | .83 | **28.36** | **15.34**<br>**41.38** | **<.01** | **10.66** | **5.96**<br>**15.36** | **<.01** |
|  | *Exponent* | -8.61 | -31.77<br>14.53 | .54 | -3.61 | -13.48<br>6.25 | .71 | .74 | -2.81<br>4.31 | .71 |
|  | *Age* | **-.65** | **-1.35**<br>**.04** | **.02** | .009 | -.28<br>.30 | .75 | -.01 | -.12<br>.09 | .75 |
|  | *Exponent x Age* | .38 | -.16<br>.93 | .14 | .01 | -.21<br>.25 | .89 | -.005 | -.09<br>.07 | .89 |
| Occipital | *Intercept* | -7.85 | -29.20<br>13.49 | .47 | **25.21** | **16.28**<br>**34.15** | **<.01** | **9.56** | **6.32**<br>**12.79** | **<.01** |
|  | *Exponent* | .40 | -17.23<br>18.04 | .97 | -1.31 | -8.69<br>6.06 | .99 | 1.72 | -.94<br>4.39 | .79 |
|  | *Age* | -.40 | -.94<br>.13 | .16 | .11 | -.11<br>.33 | .32 | -.003 | -.08<br>.07 | .89 |
|  | *Exponent x Age* | .18 | -.27<br>.64 | .56 | -.06 | -.25<br>.12 | .56 | -.01 | -.08<br>-.05 | .69 |
| Midline | *Intercept* | -8.01 | -59.99<br>43.96 | .76 | **31.53** | **9.20**<br>**53.87** | **<.01** | **10.48** | **2.38**<br>**18.58** | **.01** |
|  | *Exponent* | -.45 | -34.85<br>33.94 | .90 | -5.14 | -19.92<br>9.63 | .91 | .75 | -6.61<br>6.11 | .90 |

|  |  |  |  |  |  |  |  |  |  |
| --- | --- | --- | --- | --- | --- | --- | --- | --- | --- |
| *Age* | -.75 | -1.96 | .19 | .02 | -.49 | .66 | -.02 | -.21 | .67 |
|  |  | .46 |  |  | .54 |  |  | .16 |  |
| *Exponent x Age* | .40 | -.41 | .37 | .004 | -.35 | .84 | .001 | -.12 | .84 |
|  |  | 1.23 |  |  | .35 |  |  | .13 |  |

**Table 7.**
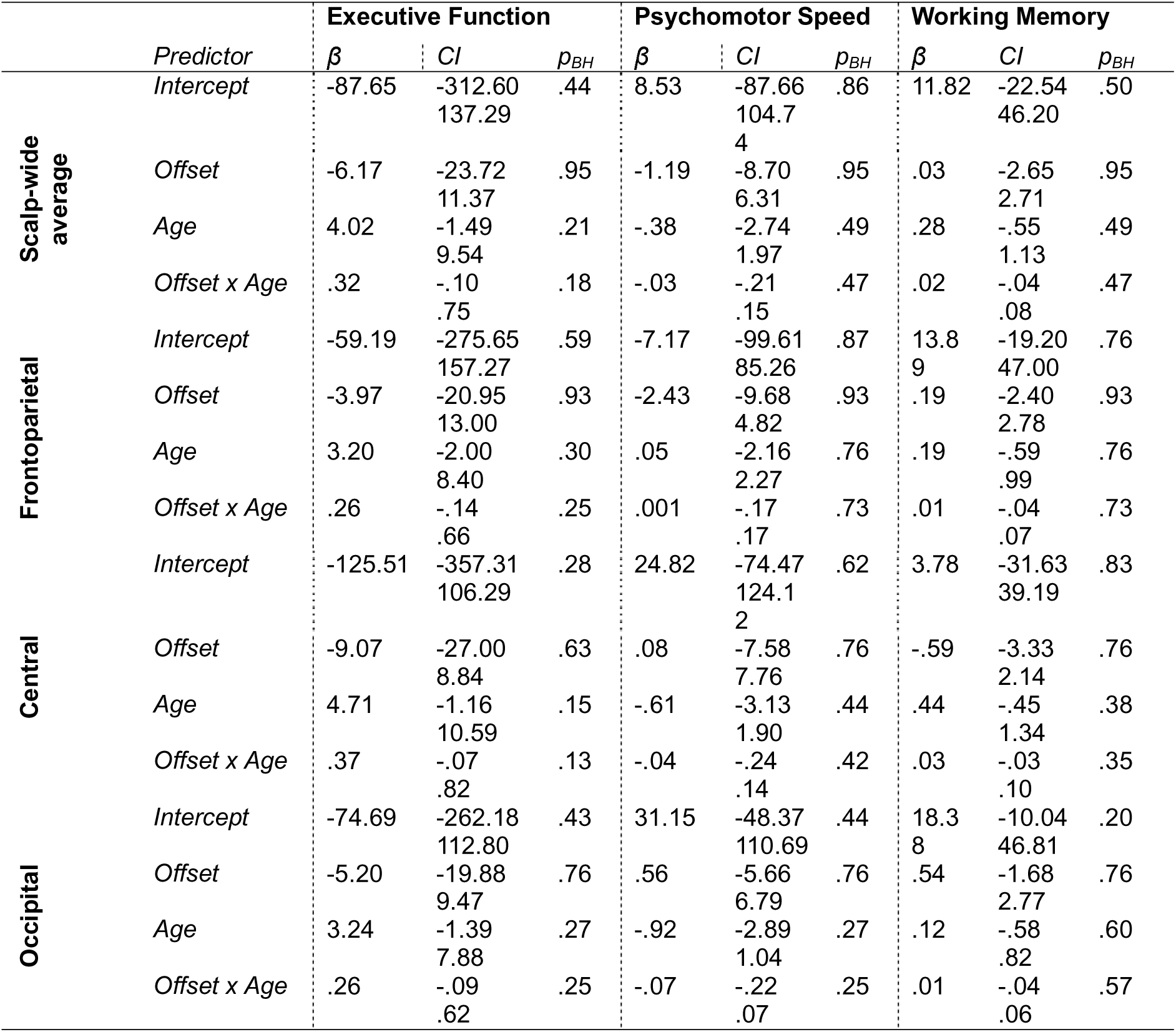
Offset by age robust regression models for cognitive domains.

|  | *Predictor* | **Executive Function** |  |  | **Psychomotor Speed** |  |  | **Working Memory** |  |  |
| --- | --- | --- | --- | --- | --- | --- | --- | --- | --- | --- |
| | | $\beta$ | *CI* | $p_{BH}$ | $\beta$ | *CI* | $p_{BH}$ | $\beta$ | *CI* | $p_{BH}$ |
| **Scalp-wide average** | *Intercept* | -87.65 | -312.60<br>137.29 | .44 | 8.53 | -87.66<br>104.74 | .86 | 11.82 | -22.54<br>46.20 | .50 |
|  | *Offset* | -6.17 | -23.72<br>11.37 | .95 | -1.19 | -8.70<br>6.31 | .95 | .03 | -2.65<br>2.71 | .95 |
|  | *Age* | 4.02 | -1.49<br>9.54 | .21 | -.38 | -2.74<br>1.97 | .49 | .28 | -.55<br>1.13 | .49 |
|  | *Offset x Age* | .32 | -.10<br>.75 | .18 | -.03 | -.21<br>.15 | .47 | .02 | -.04<br>.08 | .47 |
| **Frontoparietal** | *Intercept* | -59.19 | -275.65<br>157.27 | .59 | -7.17 | -99.61<br>85.26 | .87 | 13.89 | -19.20<br>47.00 | .76 |
|  | *Offset* | -3.97 | -20.95<br>13.00 | .93 | -2.43 | -9.68<br>4.82 | .93 | .19 | -2.40<br>2.78 | .93 |
|  | *Age* | 3.20 | -2.00<br>8.40 | .30 | .05 | -2.16<br>2.27 | .76 | .19 | -.59<br>.99 | .76 |
|  | *Offset x Age* | .26 | -.14<br>.66 | .25 | .001 | -.17<br>.17 | .73 | .01 | -.04<br>.07 | .73 |
| **Central** | *Intercept* | -125.51 | -357.31<br>106.29 | .28 | 24.82 | -74.47<br>124.12 | .62 | 3.78 | -31.63<br>39.19 | .83 |
|  | *Offset* | -9.07 | -27.00<br>8.84 | .63 | .08 | -7.58<br>7.76 | .76 | -.59 | -3.33<br>2.14 | .76 |
|  | *Age* | 4.71 | -1.16<br>10.59 | .15 | -.61 | -3.13<br>1.90 | .44 | .44 | -.45<br>1.34 | .38 |
|  | *Offset x Age* | .37 | -.07<br>.82 | .13 | -.04 | -.24<br>.14 | .42 | .03 | -.03<br>.10 | .35 |
| **Occipital** | *Intercept* | -74.69 | -262.18<br>112.80 | .43 | 31.15 | -48.37<br>110.69 | .44 | 18.38 | -10.04<br>46.81 | .20 |
|  | *Offset* | -5.20 | -19.88<br>9.47 | .76 | .56 | -5.66<br>6.79 | .76 | .54 | -1.68<br>2.77 | .76 |
|  | *Age* | 3.24 | -1.39<br>7.88 | .27 | -.92 | -2.89<br>1.04 | .27 | .12 | -.58<br>.82 | .60 |
|  | *Offset x Age* | .26 | -.09<br>.62 | .25 | -.07 | -.22<br>.07 | .25 | .01 | -.04<br>.06 | .57 |

## DISCUSSION

This study identified how resting aperiodic activity is associated with cognitive function across adulthood. Poorer performance during the Colour-Word Interference Test and Verbal Fluency Test was associated with lower exponents scalp-wide and in the frontoparietal, occipital, and midline electrode clusters. Poorer performance during the Verbal Fluency Test was also associated with smaller offsets scalp-wide and in the frontoparietal and occipital electrode clusters. Interestingly, these offset associations are moderated by age, such that better verbal fluency is associated with greater offsets in adults from 33 years of age, and this relationship strengthens across adulthood.

Increasing age was associated, to a varying degree, with poorer performance in the battery of cognitive tests consistent with Salthouse (2017). Age was also associated with lower exponents and smaller offsets, indicating greater E:I balance and neural spiking rates in younger adults. Previous studies have demonstrated that cognitive decline throughout the lifespan occurs concurrently with changes in aperiodic activity (Finley et al., 2024; Merkin et al., 2023; Thuwal et al., 2021; Tran et al., 2020; Voytek et al., 2015). Although age predicts both cognitive performance and aperiodic activity in the current study, partial correlations reveal that changes in aperiodic activity contribute to variation in verbal fluency performance and constructs of executive functioning, psychomotor speed, and working memory, independent of age. Indeed, greater scalp-wide offsets were associated with better verbal fluency scores, whereas greater exponents were associated with better performance in the Executive Function, Psychomotor Speed, and Working Memory constructs. Furthermore, the regression analyses suggest the association between the aperiodic offset and verbal fluency strengthens with age, suggesting that the influence of aperiodic activity on cognitive performance becomes more pronounced across adulthood (Voytek et al., 2015). Previous studies suggest that changes in cognitive function with aging may be attributable to a generalised factor that is shared across many domains of cognition (Salthouse, 2017). Our robust regressions suggest that an intricate relationship between age, aperiodic activity, and cognitive performance exists.

### Associations between aperiodic activity and verbal fluency

Many cognitive tasks require higher-order executive functioning that integrates lower-order cognitive processes (Berg et al., 2016; Kaplan, 1990). Aging progressively impairs executive functioning due to anatomical changes to neural circuits and depletion of neurotransmitters. Of note, the frontal-striatal network of the prefrontal cortex, integral for verbal fluency (Ghanavati et al., 2019), is particularly susceptible to age-related changes that can affect cognitive function (Buckner, 2004). This can result in difficulty performing cognitively demanding tasks as aging progresses, including the Verbal Fluency Test (McDowd et al., 2011).

The Verbal Fluency Test discriminates between healthy aging and mild cognitive impairment (McDonnell et al., 2020; Sebaldt et al., 2009). Previously, verbal fluency has been demonstrated to remain stable or increase from early to mid-adulthood then declining around 60 years of age (Whitley et al., 2016) but has also been demonstrated to remain relatively consistent in later life (50 - 75 years; Clark et al., 2009; Price et al., 2012). Younger adults outperform older adults in verbal performance consistently, which may be due to changes in the prefrontal cortex (Kim et al., 2021). Indeed, under-recruitment of the prefrontal cortex has been seen in older adults compared to their younger counterparts, which coincide with reductions in verbal memory and retrieval performance (Kapur et al., 1996; Logan et al., 2002). This has been suggested to be due to a reduction in neural plasticity with age and an inability to reorganise neurocognitive networks (Cabeza et al., 2002).

Early investigations into EEG and verbal fluency across the lifespan have demonstrated that theta activity is important for verbal fluency performance (Brickman et al., 2005; Mousavi et al., 2020). Prior studies have identified that resting-state theta activity is conflated by aperiodic activity (Finley et al., 2022). Although parameterized theta activity was not quantified in the current study due to a poor retainment of theta activity once accounting for underlying aperiodic activity, it may in fact be aperiodic activity and not theta activity that is associated with the preservation of verbal fluency with aging. The measurement of (non-parameterized) oscillatory activity is particularly susceptible to being conflated with aperiodic activity. This is because individual variation in the aperiodic slope can induce variations in the patterns of activity across all frequencies, which accounts for the majority of variation observed in total oscillatory power (Donoghue, Dominguez, et al., 2020; Donoghue et al., 2022). Therefore, there is contention regarding the presence of theta activity and to what extent changes in theta activity can be accounted to underlying changes in aperiodic activity, with new evidence suggesting differences in resting theta activity are likely due to aperiodic activity (Cesnaite et al., 2023; Finley et al., 2022; McKeown et al., 2023). However, if aperiodic activity conflates task-related theta activity is still a point of debate. As the current study assessed aperiodic activity during resting EEG recordings, the associations of aperiodic activity and cognitive performance remains during task-related EEG recordings warrants further investigation.

We determined that smaller offsets at frontoparietal, central, and occipital clusters and scalp-wide averages, were associated with poorer performance on the Verbal Fluency Test. What we did not initially expect was that these associations were present in adults as early as 33 years of age (Clark et al., 2009; Price et al., 2012; Whitley et al., 2016). As smaller offsets indicate a decrease in neural spiking rate (Donoghue, Haller, et al., 2020; Gao et al., 2017), the presence of poorer performance during the Verbal Fluency Test indicates that maintenance of neural spiking from early adulthood onwards may be beneficial for maintaining higher-order cognitive processing throughout the lifespan.

### Considerations

Investigations into the associations between aperiodic EEG activity and cognition are still in their infancy, however, this is not the first study to adopt a similar methodology (Euler et al., 2024). In the current study, we identified correlations between the aperiodic exponent and the constructs Executive Function, Psychomotor Speed, and Working Memory. However, these correlations were not seen for the aperiodic offset, nor did this result in main or interaction effects in the cognitive domain models. This is surprising considering our single assessments of each neuropsychological test identified that greater aperiodic offset was significantly associated with verbal fluency performance, the measure of which was included in the construct of Executive Function. Euler et al. (2024) recently investigated how electrode clusters of activity associate with “domains” of cognition (i.e., working memory, perceptual reasoning, processing speed, and verbal comprehension). However, there were no associations of the exponent with any specific cognitive process, but rather the scalp-wide average was associated with a general intelligence *g* factor in a cohort of 165 adults (18 – 52 years). These findings support the notion of a common influencing factor contributing to differences in cognitive performance with age (Salthouse, 2017). However, as age impacts aperiodic activity and cognition substantially in the later years of life, the inclusion of a cohort in the current study containing a larger age range may explain why aperiodic activity and age interactions were found for specific neuropsychological tests, such as verbal fluency, and not broad cognitive domains.

Factors beyond aging can impact an individual’s cognitive performance that were not investigated in the current study. One such factor is education, which has been shown to be a significant predictor of cognitive performance throughout the lifespan (Lövdén et al., 2020). Recently, slower processing times and lower working memory capability was associated with greater exponents in older adults with a high level of education but this relationship was not seen in individuals with a low level of education (Montemurro et al., 2024). Although not reported in the demographic information for the current dataset, it is possible there are differences in education level between the younger and older adults in the cohort. Education level should be considered in future studies and should be controlled for and examined as a potential moderator.

Using resting state EEG to assess brain activity provides a high temporal resolution analysis of brain activity over time; however, it is limited in its spatial sensitivity and ability to capture task-related EEG changes when recorded simultaneously during a task. Although we have identified that reductions in resting state EEG aperiodic activity coincide with worse verbal fluency performance, we are unable to determine which neural structures are responsible for this association. Functional MRI studies have identified that verbal fluency performance is particularly reliant on the function of prefrontal, anterior cingulate gyrus, and entorhinal areas (De Marco & Venneri, 2021; Schaufelberger et al., 2005). It may be the case that these changes in the activity of these areas are responsible for the determinants seen in the current study. However, as reductions in aperiodic EEG activity occurred scalp-wide, it may be that broad reductions in aperiodic activity, likely reflecting changes in the dynamics of postsynaptic potentials (Gao et al., 2017; Miller et al., 2009), across multiple brain regions are responsible for reductions in verbal fluency performance with age.

### Conclusion

Our study examined variations in resting aperiodic EEG activity with age and how they relate to cognitive performance. This comprehensive approach expands our understanding of the cognitive implications of aperiodic activity. Aperiodic activity was associated with performance during the Colour-Word Interference Test and Verbal Fluency Test. Aperiodic activity and age interactions were observed for the Verbal Fluency Test, where smaller offsets were associated with progressively worse Verbal Fluency Test performance with age, with associations appearing as early as 33 years of age. The presence of alterations in aperiodic activity and cognitive performance suggests that aperiodic activity may be an increasingly significant factor that may contribute to the preservation of an individual’s cognitive function earlier than previously understood, even as the relationship becomes more pronounced with age.

## Abbreviations

CWIT: Colour-Word Interference Test
DKEFS: Delis-Kaplan Executive Function System score
E:I: Excitation: inhibition
FFT: Fast Fourier Transformation
IAPF: Individual alpha peak frequency
SRM: Stimulus-Selective Response Modulation
OLS: Ordinary least squares
PCA: Principal components analysis
PSD: Power spectral density
RAVLT: Reys Adult Verbal Learning Test
SOBI: Second order blind identification
WAIS-IV DST: Wechsler Adult Intelligence Scale IV Digit Span Test

## ACKNOWLEDGEMTENTS

The authors would like to acknowledge the authors of the SRM Resting-state EEG dataset – Christoffer Hatlestad-Hall, Trine Waage Rygvold, and Stein Andersson

